# Electrophysiological recordings reveal photoreceptor coupling in the dorsal rim areas of honeybee and bumblebee eyes

**DOI:** 10.1101/2025.04.29.651169

**Authors:** George E Kolyfetis, Gregor Belušič, James J Foster

## Abstract

Many insects rely on skylight polarization patterns to navigate their habitats. To perform this vital task, most insect species have evolved specialized ommatidia in the dorsal rim area (DRA) of their compound eyes that are adapted to detect linearly polarized light in large patches of sky. In this study, we conducted electrophysiological recordings of honeybee (*Apis mellifera*) and bumblebee (*Bombus terrestris*) photoreceptors in the DRA and other regions of the compound eye to map their receptive fields. For both species, we report novel evidence for photoreceptor coupling, i.e., spatial summation, present in the retinal layer of the DRA. We explore spatial summation as a possible, eye-region-specific mechanism to increase the effective size of DRA ommatidia receptive fields; a crucial functional feature of the polarization compass.

## Introduction

Light arriving from the sun gets scattered and polarized in the earth’s atmosphere, creating patterns of polarization in the sky (*1, 2*). These patterns are dependent on the sun’s position and can thus act as a valuable navigational reference system for the many insects that perceive them (*2–4*). Honeybees have been behaviourally demonstrated to utilize an internal map of skylight polarization’s topography, a ‘polarization compass’, to estimate the position of the sun and navigate successfully (*5, 6*).

Most insect species detect polarized light using specialized, polarization-sensitive photoreceptors in the Dorsal Rim Area (DRA) of their compound eyes (*7*). In honeybees and bumblebees, these are UV-sensitive photoreceptors with high polarization sensitivity (PS) (*7– 10*). Honeybee DRA photoreceptors are also known to possess exceptionally wide receptive fields (RFs) compared to main retina and dorsal-eye photoreceptors (*7*). The shape of these wide RFs has been previously described in honeybees; they exhibit a central region with high relative sensitivity and a wide ‘brim region’ surrounding the centre, where sensitivity decreases very gradually with increasing off-axis angle (*7*).

Little is known about the mechanism that underlies the wide RFs of the DRA ommatidia. It has been previously proposed that light-scattering structures, such as rugged-walled pore canals, in the corneal lenses of the DRA ommatidia of several Hymenopteran species (*11*) could potentially result in increased sensitivity to off-axis illumination, and thus extended RFs (*7, 12*). The existence of pore canals or wide RFs in the DRA of bumblebees remains unknown. For crickets, both anatomical (lack of screening pigment and corneal facets) (*13*) and neural (spatial integration via polarization-sensitive interneurons) (*14, 15*) adaptations have been proposed as the source of wide DRA receptive fields. Neural spatial integration of this type, acting as a spatial low pass-filter, can smooth out local disturbances in the skylight polarization pattern (e.g., due to clouds), increase the polarization signal-to-noise ratio and thus aid navigational efficiency (*14*). Even though wide RFs naturally result in only low spatial resolution, DRA photoreceptors have been reported to have high absolute sensitivity, i.e., capture a high proportion of available photons (*13, 16*).

In this study, we mapped the receptive fields (RFs) of honeybee (*Apis mellifera*) and bumblebee (*Bombus terrestris*) dorsal rim area (DRA) photoreceptors by recording from them electrophysiologically. We identified photoreceptor coupling at the retinal layer of both species’ DRA, which constitutes novel evidence of eye-region-specific neural spatial summation present at the retinal layer. We discuss cell coupling as a potential, plastic mechanism for increasing the effective size of bee DRA receptive fields, and, in light of this new discovery, we reexamine the function of pore canals in polarization vision.

## Methods

### Electrophysiological Photoreceptor Recordings

We immobilized honeybee (*Apis mellifera mellifera* and *Apis m. carnica*) and bumblebee (*Bombus terrestris dalmatinus*) workers in plastic pipette tips using beeswax and resin and carefully aligned the head with the stimulus. For both main retina and DRA recordings, a microelectrode (0.5/1.0 mm inner/outer diameter borosilicate glass, filled with 3M KCl, resistance R > 100 MΩ) was inserted into the cornea ventral to the DRA, through a small hole (Fig. 1a), connected to an amplifier (SEC10LX, NPI electronic, Germany) and advanced with a piezo micromanipulator (Sensapex, Finland). The experiments were conducted in a dark laboratory, in a Faraday cage made of iron plates, using dense stimulation with bright light flashes, so that the animals were not dark-adapted.

**Fig. 1.**
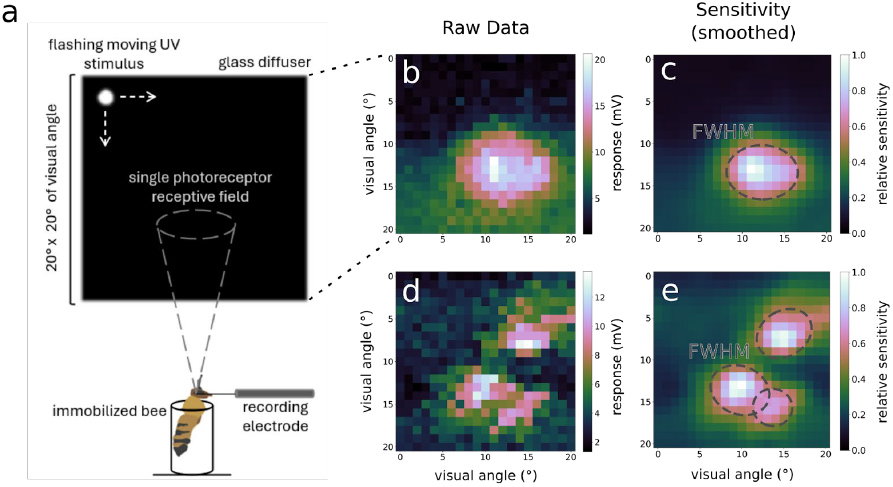
Experimental setup and receptive field mapping. (a) Overview of the experimental setup. Bees were immobilized and presented with a flashing, moving UV-stimulus while we recorded responses from single photoreceptors. (b) Raw mV and sensitivity-transformed (smoothed) (c) response output of a non-coupled photoreceptor. (d) Raw mV and sensitivity- transformed (smoothed) (e) response output of a coupled DRA photoreceptor. Dotted lines outline the Full Width at Half Maximum (FWHM) of the receptive fields. These spatial sensitivity recordings allowed us to map individual and coupled receptive fields.

All cells were initially stimulated with a multispectral light synthesiser (LED synth) (*17*) for 2 seconds to assess their spectral sensitivity. We fitted a visual pigment template function to the spectral responses (*18*). The polarization sensitivity (PS) of the photoreceptors was evaluated with a flash stimulus from a Xe arc lamp and a monochromator (B&M Optik, Germany), projected through a polarizer (OUV2500, Knight Optical, UK) at the peak wavelength of the determined spectral sensitivity. Then, unpolarized flashes at the same wavelength, projected through a graded ND filter (max. intensity in the UV ∼10^14^ photons cm^-2^ s^-1^), were used to measure the intensity-response function (“ V-log(I)”). The single cell responses were non-linearly transformed into sensitivity values using the cell’s intensity response function and an inverse Hill transformation (*19*). We then calculated polarization sensitivity as the ratio between maximum and minimum sensitivity to light, polarized at different angles (*20*). Where multiple polarization response measurements were made for a single cell, a per-angle arithmetic mean of the raw measurements was calculated first. Mean PS values above 20 were considered artifactual owing to the low or even hyperpolarizing minimum response to polarized light measured (possibly due to inhibition from polarization-opponent photoreceptors) (*21*) and were thus excluded.

We mapped the spatial receptive fields of UV-sensitive cells by presenting flashes from a 365 nm UV LED (Cairn, UK) at different positions on a rear-illuminated, UV-transmissive glass diffuser using a pair of AT20L-L galvanometer mirrors (Shenzhen City Aoxinjie Technology, China). The centre of the photoreceptor receptive field was determined by using moving horizontal and vertical bars and the whole receptive field was mapped with a bright, flashing 0.9° x 0.9° square stimulus, moving left to right, top to bottom on a 20° x 20° grid (Fig. 1a). PS was re-measured in some DRA cells which showed coupling, by aligning their multiple optical axes with the stimulus and measuring PS in the different parts of their receptive field. For more information on the photoreceptor recordings see the Supplementary Methods.

### Receptive Field Modelling

To model the receptive fields of the main retina and DRA UV-sensitive cells (16 honeybee cells and 31 bumblebee cells in total), we divided every row of the 20° x 20° raw spatial sensitivity (SS) data grid in 21 equal parts and extracted the maximum voltage response values for each. From the maximum values, we subtracted the minimum response of each part, which roughly corresponds to the background noise. These modified raw SS data (Fig. 1b,d) were smoothed using Gaussian filtering (scipy.ndimage.gaussian_filter; sigma=1°, all other parameters set to default) to acquire the smoothed SS data. We then applied the inverse Hill function to normalized SS datasets to acquire the final SS data (Fig. 1c,e, Fig. 2).

**Fig. 2.**
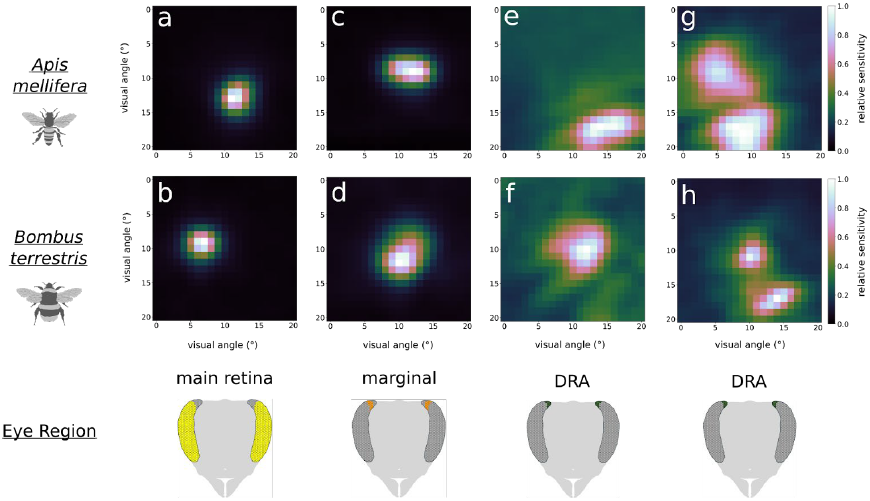
Receptive field shapes in different eye regions of honeybees and bumblebees. Main retina (a,b), marginal DRA (c,d), non-coupled DRA (e,f) and coupled DRA (g,h) photoreceptor receptive field shapes for honeybee (*Apis mellifera*) and bumblebee (*Bombus terrestris*) respectively. In both species, receptive field size tends to increase from the main retina to the DRA, and their shapes become more irregular. Coupled DRA photoreceptor receptive fields were observed in both species.

The size of the photoreceptor receptive fields was quantified using the fitted 2D, elliptical Gaussian fields. For coupled photoreceptors, we added a ‘weighting’ parameter to each of the RFs to distinguish the main (punctured; high weighting) cell from the coupled one(s) (distant from the electrode tip; low weighting). For all summary plots, only the properties of the main cells were considered. Recorded cells were categorized into main retina, marginal DRA and DRA based mainly on the recording depth (topologically), but also their intensity response dynamics (main retina and marginal DRA: high sensitivity; DRA: low sensitivity), polarization sensitivity (main retina: low PS, marginal and DRA: high PS) and approximate size of their receptive fields.

For more information on the RF modelling see the Supplementary Methods.

### Statistical Analyses and Visualizations

A 2-way ANOVA was performed on PS and FWHM data (Fig. 3a,c). PS and FWHM residuals were found to be normally distributed (Shapiro-Wilk p = 0.2093 and p = 0.9255 respectively). Tukey’s HSD post hoc tests were carried out on both PS and FWHM data.

**Fig. 3.**
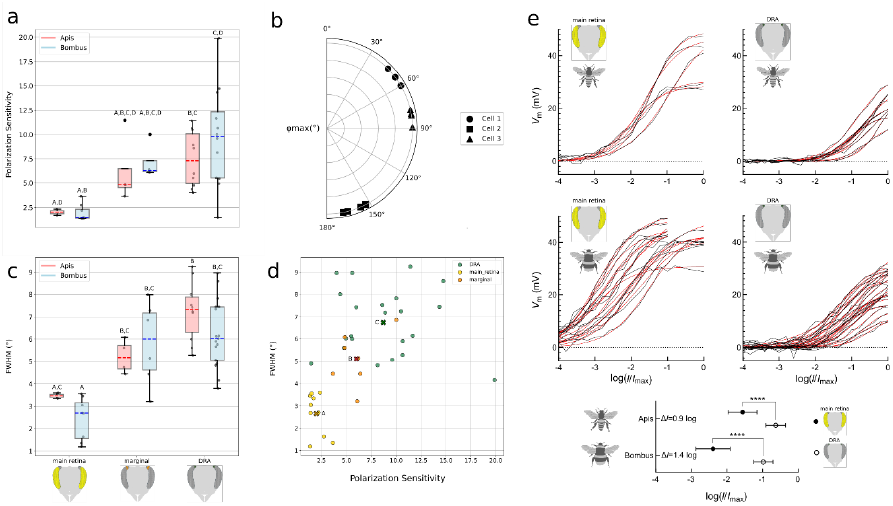
Overview of visual properties in the different eye regions. (a) Polarization Sensitivity in the different eye regions. (b) *φ*_max_ angle differences between the multiple receptive fields in 3 different DRA photoreceptor cells. (c) Receptive field size (represented by their FWHM) in the different eye regions. (d) Relationship between RF size (FWHM) and PS in the different eye regions. (e) Intensity-response curves of UV receptors in different eye regions (black curves) in both species; lower panel, half-maximum intensity parameters of Hill functions (red curves), fitted to voltage data. Whiskers in (a) and (b) represent Q1 − 1.5×IQR (lower whisker) and Q3 + 1.5×IQR (upper whisker). Different letters indicate significant differences.

A linear model accounting for differences between eye regions and species was fitted to the FWHM∼PS data (see Supplementary Table 1) after a model selection process. Residuals were normally distributed (Shapiro-Wilk p = 0.5282), and a Tukey’s HSD post hoc test was performed.

All statistical analyses were performed in Python using the scipy and statsmodels packages. All Figure visualizations were carried out using Python’s Matplotlib (*22*) and Inkscape (*23*).

## Results

We performed electrophysiological recordings of 17 honeybee (*Apis mellifera*) and 31 bumblebee (*Bombus terrestris*) UV-sensitive photoreceptors in the main retina (n=10), marginal DRA (n=11) and DRA (n=27) to test their spatial, spectral and polarization sensitivity. We also recorded from blue-sensitive photoreceptors in the main retina, but they were excluded from further analyses. Honeybees’ and bumblebees’ peak wavelength sensitivity of UV photoreceptors lay at ∼340 nm, similar to previous reports (*7, 10*).

DRA photoreceptors have the largest RFs across the different eye regions in both honeybees and bumblebees (Fig. 2, Fig. 3c). Furthermore, in 3 honeybee and 3 bumblebee spatial sensitivity recordings, DRA cells exhibited multiple regions of high sensitivity (>0.5) that were separated by regions of low sensitivity (usually < 0.2). These RFs likely belong to neighbouring, coupled photoreceptor cells which that only partly overlap in their visual fields, but share their responses. Examples of this photoreceptor coupling in the DRA can be seen in Fig. 1d,e and Fig. 2g,h. Analysis of response delay differences and PS angular maxima between primary and secondary RFs confirmed that the multiple RFs were not the result of optical artefacts (see Supplementary Material).

The difference in RF size between ommatidia in different eye regions can be seen in Fig. 2. Main retina RFs appear to be small and circular, resembling a radially symmetric Gaussian. The sensitivity is high at the centre of the RFs but decreases rapidly at distances of 1°–2°. Marginal DRA RFs exhibit similar characteristics but have a more elliptical shape and are slightly larger compared to the main retina. DRA photoreceptors’ RFs cover the largest visual angle. They usually have irregular shapes and can be characterized by wide regions of low, non-zero sensitivity around the centres of the RFs. Their spatial sensitivity decreases more gradually from the centre of the RFs than the sensitivity of marginal and main retina cells (Fig. 1b-e, Fig. 2e-h). Coupled photoreceptors cover an even wider area of the total visual scene, combining RFs of different cells.

An overview of UV photoreceptor properties in the different eye regions can be seen in Fig. 3. Apis DRA mean PS (7.48) was significantly higher than Apis mean main retina PS (2.02) (p = 0.0222). Bombus DRA mean PS (9.71) was also significantly higher than Bombus mean main retina PS (1.95) (p = 0.0003). For marginal DRA cells, a mean PS of 6.19 was determined for honeybees and 7.18 for bumblebees (Fig. 3a).

The UV and blue receptors in the main retina had their angle of maximal PS (*φ*_max_) aligned with the dorso-ventral eye axis. The UV receptors in the DRA had their maxima loosely arranged parallel or perpendicular to the eye’s edge. In coupled DRA receptors, the *φ*_max_ of the coupled units was offset by 3–5° from the *φ*_max_ of the main unit (Fig. 3b, Supplementary Fig. 3), indicating coupling between homologous cells, arranged in a fan shape, typical for the DRA (*24*).

For ease of comparison, we transformed elliptically shaped Gaussian RFs to circular ones with equal surface area and we report their FWHM. Apis DRA mean RF (7.24°) was significantly larger than Apis and Bombus mean main retina RF (3.47°, 2.45°; p = 0.0113 and p = 0.0 respectively), Bombus DRA mean RF (6.23°) was significantly larger than Bombus mean main retina RF (p = 0.0). Furthermore, Bombus mean main retina RF was significantly smaller than Apis and Bombus marginal DRA RF (5.22°, 5.82°; p = 0.0226 and p = 6e-4 respectively) (Fig. 2, Fig. 3c).

Cells with higher PS also had larger receptive fields, both of which also differed between eye regions (Fig. 3d). After fitting a linear model accounting for species and eye region to the FWHM–PS data (Supplementary Table 1), neither PS (β = 0.170°/PS, p = 0.799) nor species (β = -0.8793°, p = 0.062) were found to significantly predict FWHM size. The model revealed significant differences between main retina RF size and marginal DRA (β = 2.1868°, p = 0.004) and DRA (β = 3.7542°, p = 0.0) and a post hoc Tukey’s HSD test confirmed the differences. Main retina cells had significantly smaller RF size than both marginal DRA and DRA (mean diff = 2.46°, p = 0.0018 and mean diff = 4.09°, p = 0.0, respectively) and marginal DRA cells had significantly smaller RF size than DRA cells (mean diff = 1.64, p = 0.0198).

The cells also had strikingly different absolute light sensitivities: the UV receptors in the main retina and marginal DRA were ∼10 times more sensitive (0.9 log units in honeybees, 1.4 log units in bumblebees) than their counterparts in the DRA (Fig. 3e).

## Discussion

In this study we identify a novel form of retina-level spatial summation, exclusive to the dorsal rim area and adjacent marginal region. While our findings otherwise confirm the previously reported properties of honeybee (*Apis mellifera*) and bumblebee (*Bombus terrestris*) DRA photoreceptors, this discovery reveals a potentially unique solution to the trade-off between polarization sensitivity and absolute sensitivity (*25*). Around a quarter of our single-cell recordings in the DRA in both species showed multiple high-sensitivity regions separated by areas of low sensitivity, suggesting photoreceptor coupling. This could act as a flexible mechanism of neural spatial summation that can be varied depending on the visual environment (*26*), allowing the visual system to balance the increase in sensitivity against the decrease in resolving power; a trade-off inherent in pooling signals from neighbouring units (*25*). Indeed, photoreceptor coupling was not observed in our recordings from the main retina high-acuity zones adapted for object vision (*33*), where low spatial resolution would be maladaptive. Although differential resolution and sensitivity across apposition eye regions is possible through varying optical parameters (*34*), a neural mechanism permits rapid activation at low resource-cost, requiring no extra cornea material. Numerous species enhance sensitivity under low-light or low-contrast conditions via neural spatial summation in the peripheral visual system, as found in the ganglion and bipolar cells in the vertebrate retina (*27–29*) and laminar monopolar cells in hawkmoths (*30*). In the *Drosophila* DRA, cell coupling takes place in the medulla where ∼10 photoreceptors converge to 1 Distal medulla DRA neuron (Dm-DRA) (*31*). However, photoreceptor coupling in the retina of the insect DRA has not been reported before. Since the primary function of the DRA is to distinguish changes in the polarization pattern in large sky patches (*6, 32*), photoreceptor coupling could eliminate cloud noise by smoothing out the polarization pattern, as proposed for polarization-sensitive interneuron coupling in the cricket DRA (*15*).

We find that UV-sensitive DRA photoreceptors in both bee species possess wide RFs and high PS. These large, overlapping RFs would capture a wide sky area, likely increasing signal and making navigation more efficient (*7, 14*). Rugged-walled pore canals in the DRA have been proposed to widen the ommatidial RFs by scattering incoming light (*7, 11*). Although we do not present any direct evidence in favour of this phenomenon here, we propose spatial summation via photoreceptor coupling as another cause of the wide RFs of the DRA. Pore canals could act as an aperture-limiter for the DRA ommatidia, scattering a portion of incident light out of the light path to reduce the quantity that reaches the rhabdoms. Since DRA ommatidia usually view the bright sky, this mechanism could prevent the otherwise rapid saturation of their photoreceptors. Indeed, the DRA intensity-response curves were steeper than in the main retina (Fig. 3e) but showed lower sensitivity. This would allow the DRA receptors to better encode the small contrast changes that would be induced by changes in skylight polarization, while avoiding saturation at typical skylight intensities. Spatial summation via photoreceptor coupling may help compensate for this low sensitivity under low-contrast conditions. Since our stimuli were bright, sub-saturating flashes, the photoreceptors were not dark-adapted, it appears that this spatial summation mechanism is active under daylight conditions.

The observed DRA photoreceptor coupling could also potentially explain the wide low-sensitivity areas of the spatial sensitivity curve (*7*) previously attributed to the pore canals (12). Contributions from coupled photoreceptors’ RFs would have been difficult to identify in previous studies, since the stimuli presented spanned only two orthogonal axes. However, it is also possible that the two mechanisms act complementarily, pore canals widening each ommatidium’s RF and cell coupling expanding each photoreceptor’s effective RF (*7*) (Fig. 2e- h). The wide low-sensitivity areas (brims) of the spatial sensitivity curve described in Labhart 1980 may be captured by the ‘offset’ parameter of our Gaussian models of the DRA RFs (see Methods). Future work focused on the function of the pore canals will be crucial to elucidate their role in insect polarization vision.

## Conclusion

The results of this work provide the first evidence for the coupling of photoreceptors between adjacent rhabdoms in the dorsal rim areas (DRA) of honeybee (*Apis mellifera*) and bumblebee (*Bombus terrestris*) compound eyes. Our electrophysiological recordings show that DRA photoreceptors exhibit multiple receptive fields separated by regions of low spatial sensitivity. Slight changes in φ_max_ angles as well as response delay differences further corroborate the existence of photoreceptor coupling in the DRA. Importantly, spatial summation in the DRA may serve a crucial functional role, reducing noise in the detection of skylight polarization, and thus aiding navigation in bees. Overall, this study contributes valuable insights into bees’ visual systems and adaptations of the DRA for polarization vision.

## Ethics

This work did not require ethical approval from a human subject or animal welfare committee.

## Data accessibility

All Data files needed to evaluate the conclusions in the paper have been uploaded to Figshare under: https://doi.org/10.6084/m9.figshare.28890938.v1 Custom scripts used for analyses, statistical tests and visualizations can be found in the GitHub repository: https://github.com/JJFosterLab/DRA-coupling

## Declaration of AI use

We have not used AI-assisted technologies in creating this article.

## Authors’ contributions

G.E.K.: conceptualization, data curation, visualization, formal analysis, investigation, methodology, writing – original draft, writing – review and editing; G.B.: conceptualization, conduction of experiments, investigation, methodology, visualization, writing – review and editing; J.J.F.: conceptualization, investigation, methodology, writing - review and editing.

## Conflict of interest declaration

We declare we have no competing interests.

## Funding

J.J.F. & GEK were supported by a DFG research grant awarded to J.J.F. (project number 451057640). J.J.F. & G.E.K. also received funds from the DFG Centre of Excellence “ Centre for the Advanced Study of Collective Behaviour” (EXC 2117 - 422037984). G.B. was supported by the Air Force Office of Scientific Research (grant no. FA8655-23-1-7049) and Javna Agencija za Raziskovalno Dejavnost RS (grant no. P3-0333).

## Acknowledgements

We thank Anna Stöckl for help with animals and feedback on the manuscript. We also thank Frida Hildebrandt for help with animals and Andrew McCauley for helpful suggestions for analyses. We thank Marko Ilić for helping with the experimental setup.

## Notes

https://doi.org/10.6084/m9.figshare.28890938.v1

https://github.com/JJFosterLab/DRA-coupling

### Competing Interest Statement

The authors have declared no competing interest.

## References

1. J. Hecht, Optics: light for a new age (Jeff Hecht, 1987).

2. J. Foster, G. Horváth, Ed. (Springer Nature Switzerland, Cham, 2024; 10.1007/978-3-031-62863-4_2), pp. 19–38.

3. B. Webb, The internal maps of insects. J. Exp. Biol. 222, jeb188094 (2019).

4. T. L. Warren, Y. M. Giraldo, M. H. Dickinson, Celestial navigation in Drosophila. J. Exp. Biol. 222, jeb186148 (2019).

5. M. L. Brines, J. L. Gould, Bees have rules. Science (80-.). 206, 571–573 (1979).

6. S. Rossel, R. Wehner, How bees analyse the polarization patterns in the sky: experiments and model. J. Comp. Physiol. A. 154, 607–615 (1984).

7. T. Labhart, Specialized photoreceptors at the dorsal rim of the honeybee’s compound eye: polarizational and angular sensitivity. J. Comp. Physiol. 141, 19–30 (1980).

8. R. Menzel, A. W. Snyder, Polarised light detection in the bee, Apis mellifera. J. Comp. Physiol. 88, 247–270 (1974).

9. V. B. Meyer-Rochow, Electrophysiology and histology of the eye of the bumblebee Bombus hortorum (L.)(Hymenoptera: Apidae). J. R. Soc. New Zeal. 11, 123–153 (1981).

10. P. Araújo, G. Belušič, M. Ilić, J. Foster, K. Pfeiffer, E. Baird, Polarized light detection in bumblebees varies with light intensity and is mediated by both the ocelli and compound eyes. Biol. Lett. 20, 20240299 (2024).

11. F. Aepli, T. Labhart, E. P. Meyer, Structural specializations of the cornea and retina at the dorsal rim of the compound eye in hymenopteran insects. Cell Tissue Res. 239, 19–24 (1985).

12. E. P. Meyer, T. Labhart, Pore canals in the cornea of a functionally specialized area of the honey bee’s compound eye. Cell Tissue Res. 216, 491–501 (1981).

13. T. Labhart, B. Hodel, I. Valenzuela, The physiology of the cricket’s compound eye with particular reference to the anatomically specialized dorsal rim area. J. Comp. Physiol. A. 155, 289–296 (1984).

14. T. Labhart, How polarization-sensitive interneurones of crickets see the polarization pattern of the sky: a field study with an opto-electronic model neurone. J. Exp. Biol. 202, 757–770 (1999).

15. T. Labhart, J. Petzold, H. Helbling, Spatial integration in polarization-sensitive interneurones of crickets: a survey of evidence, mechanisms and benefits. J. Exp. Biol. 204, 2423–2430 (2001).

16. D. Brunner, T. Labhart, Behavioural evidence for polarization vision in crickets. Physiol. Entomol. 12, 1–10 (1987).

17. G. Belušič, M. Ilić, A. Meglič, P. Pirih, A fast multispectral light synthesiser based on LEDs and a diffraction grating. Sci. Rep. 6, 32012 (2016).

18. D. G. Stavenga, On visual pigment templates and the spectral shape of invertebrate rhodopsins and metarhodopsins. J. Comp. Physiol. A. 196, 869–878 (2010).

19. J. Zeil, W. A. Ribi, A. Narendra, in Polarized light and polarization vision in animal sciences (Springer, 2014), pp. 41–60.

20. G. D. Bernard, R. Wehner, Functional similarities between polarization vision and color vision. Vision Res. 17, 1019–1028 (1977).

21. R. C. Hardie, Properties of photoreceptors R7 and R8 in dorsal marginal ommatidia in the compound eyes of Musca and Calliphora. J. Comp. Physiol. A. 154, 157–165 (1984).

22. J. D. Hunter, Matplotlib: A 2D graphics environment. Comput. Sci. Eng. 9, 90–95 (2007).

23. T. Bah, Inkscape: guide to a vector drawing program (prentice hall press, 2007).

24. T. Labhart, Can invertebrates see the e-vector of polarization as a separate modality of light? J. Exp. Biol. 219, 3844–3856 (2016).

25. F. J. H. Heras, S. B. Laughlin, Optimizing the use of a sensor resource for opponent polarization coding. PeerJ. 5, e2772 (2017).

26. E. J. Warrant, Seeing better at night: life style, eye design and the optimum strategy of spatial and temporal summation. Vision Res. 39, 1611–1630 (1999).

27. K. Donner, Adaptation-related changes in the spatial and temporal summation of frog retinal ganglion cells. Acta Physiol. Scand. 131, 479–487 (1987).

28. J. B. Demb, K. Zaghloul, L. Haarsma, P. Sterling, Bipolar cells contribute to nonlinear spatial summation in the brisk-transient (Y) ganglion cell in mammalian retina. J. Neurosci. 21, 7447–7454 (2001).

29. C. Stone, L. H. Pinto, Response properties of ganglion cells in the isolated mouse retina. Vis. Neurosci. 10, 31–39 (1993).

30. A. L. Stöckl, W. A. Ribi, E. J. Warrant, Adaptations for nocturnal and diurnal vision in the hawkmoth lamina. J. Comp. Neurol. 524, 160–175 (2016).

31. G. Sancer, E. Kind, H. Plazaola-Sasieta, J. Balke, T. Pham, A. Hasan, L. O. Münch, M. Courgeon, T. F. Mathejczyk, M. F. Wernet, Modality-specific circuits for skylight orientation in the fly visual system. Curr. Biol. 29, 2812–2825 (2019).

32. S. Rossel, R. Wehner, in Neurobiology and behavior of honeybees (Springer, 1987), pp. 76–93.

33. A. Kelber, H. Somanathan, Spatial vision and visually guided behavior in Apidae. Insects. 10, 418 (2019).

34. G. J. Taylor, P. Tichit, M. D. Schmidt, A. J. Bodey, C. Rau, E. Baird, Bumblebee visual allometry results in locally improved resolution and globally improved sensitivity. Elife. 8, e40613 (2019).

