## Supplementary Material for "Electrophysiological recordings reveal photoreceptor coupling in the dorsal rim areas of honeybee and bumblebee eyes"

### Supplementary Methods

##### Electrophysiological Photoreceptor Recordings

To adjust for the saturation of the photoreceptors in their intensity responses, we set the first data point after the maximum response (and all points thereafter) equal to the max response if the point was smaller than 0.95 x <max response>, as this indicated that the cell had already started to saturate. We fitted (scipy.optimize.curve_fit) a cosine-squared function to the response values of the polarization sensitivity recordings to predict more accurate maximum and minimum response amplitudes.

##### Receptive Field Modelling

We normalized the smoothed SS data with the following algorithm to remap them in the range [0, 1]: a) we divided by the predicted maximum response of the Vlog(I) sigmoid function (Emax) if the ratio <max_smoothed_values> / Emax < 1 (i.e., the maximum SS response was smaller than the maximum intensity response), b) if the ratio was between 1 and 1.05 (i.e., the maximum SS response was slightly larger than the maximum intensity response), we again normalized by the predicted Emax, but replaced all values > 1 with 1, c) if the ratio was > 1.05 (i.e., the maximum SS response was much larger than the maximum intensity response) and <max_smoothed_values> < 40mV (empirically determined average maximum intensity response), we normalized by the average Emax of all recordings that were <max_smoothed_values> < Emax < 40mV and d) if ratio > 1.05 (i.e., the maximum SS response was much larger than the maximum intensity response) and <max_smoothed_values> > 40mV, we normalized by the average Emax of all recordings that were Emax > <max_smoothed_values > 40mV.

To determine the number of receptive fields present in each recording, we used contour maps (matplotlib.pyplot.contour) with threshold levels of 0.5–0.7 relative sensitivity. This allowed us to define regions of high sensitivity separated by regions of low sensitivity, a pattern that indicates multiple receptive fields. Then, we fitted 1–3 2D, elliptical Gaussian functions to the receptive fields, the parameters of which were estimated using maximum likelihood, with Bayesian priors for receptive field centres close to the contour's centroid and similar full-width at half-maximum to those previously described for honeybees' DRAs. We added an extra 'offset' parameter to the Gaussian fits to account for the variable noise in the different recordings. For simplicity, we report the full-width at half maximum (FWHM) of circular receptive fields with equal area to the elliptical ones (Fig. 3c,d). Where we had multiple SS recordings for a single cell, an average circular FWHM (of high-quality recordings) was calculated.

##### Response timing in coupled photoreceptors

As an indicator of chemical or electrical coupling (synapses or gap junctions respectively) between photoreceptors, we used the difference in response time between the coupled cells. To that aim, we fitted a 'Delayed Normalization' (DN) response function (*1*) to our response data at the centre of the RFs (scipy.optimize.curve_fit) (Supplementary Fig. 1).

The Zhou et al. model (*1*) encompasses a 'time-to-peak' parameter ($\tau1$) which we directly compared between the coupled cells. As initial parameter values, we set the default values reported in Zhou et al., 2019 (*1*). We set the weight (w) parameter to 0 as recommended, i.e., to reduce the number of free parameters and because the effects of 'w' are negligible in most cases. Apart from $\tau1$, we also directly compared the time since the start of the flashing stimulus until the maximum predicted response (from the DN model). We also compared response delays across the whole extent of coupled cells' RFs (Supplementary Fig. 2).

### Supplementary Results

To discriminate between cell coupling (electrical via gap junctions or chemical via synapses), and optical properties of the dioptrical apparatus, we fitted a 'Delayed Normalization' curve (*1*) to our response data of coupled cells (at the centre of their RFs; Supplementary Fig. 1) to examine their times-to-peak (time since the start of the stimulus until the maximum predicted response value; $\tau1$ parameter). We identified a $\tau1$ difference of 0–5 ms between the main and secondary cells. Where there were differences, the main cells always responded faster than the secondary cells (Supplementary Fig. 1 and 2). This difference in response delay confirms that the multiple RFs are due to cell coupling and not other visual properties. The type of cell coupling could not be identified through our response delay analyses. Thus, further experiments are required to resolve this, e.g., using cell stainings.

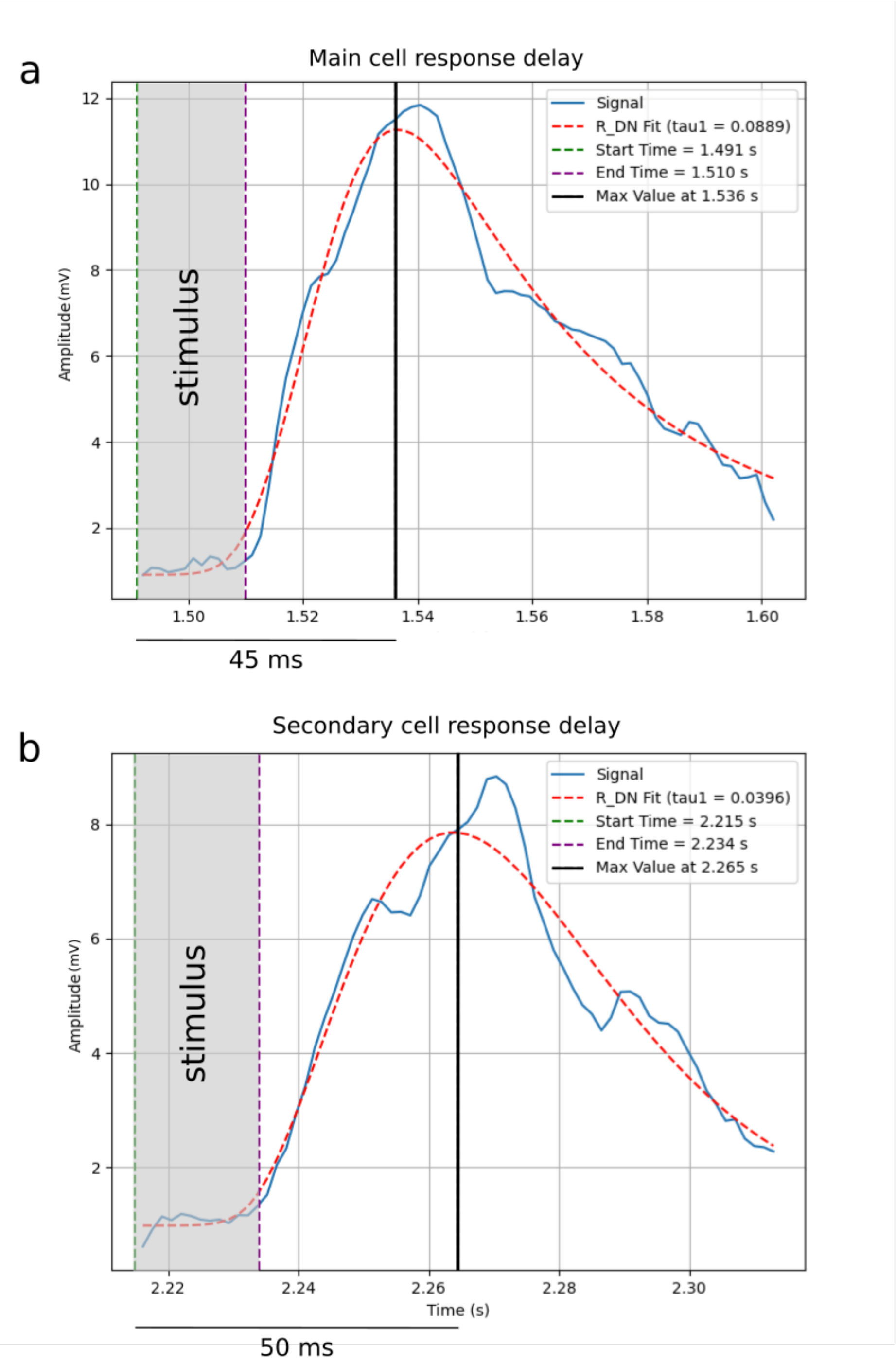

**Supplementary Fig. 1. Example of 'Delayed Normalization' (DN) model fits to coupled cells' responses.** (a) The DN fit predicts a maximum response at 45ms after stimulus start at the center of the RF. (b) The DN fit predicts a maximum response at 50ms after stimulus start at the center of the RF. The centers of the receptive fields were defined as the means (x,y) of the 2D Gaussian fits. The 'time-to-peak' (τ1) parameter of the model can be directly compared between the coupled cells, although the 'max_fitted_response_time - stimulus_start_time' is more accurate in some cases. This example corresponds to Fig. 1e and Supplementary Fig. 2d, main (low left) and secondary (upper right) RFs.

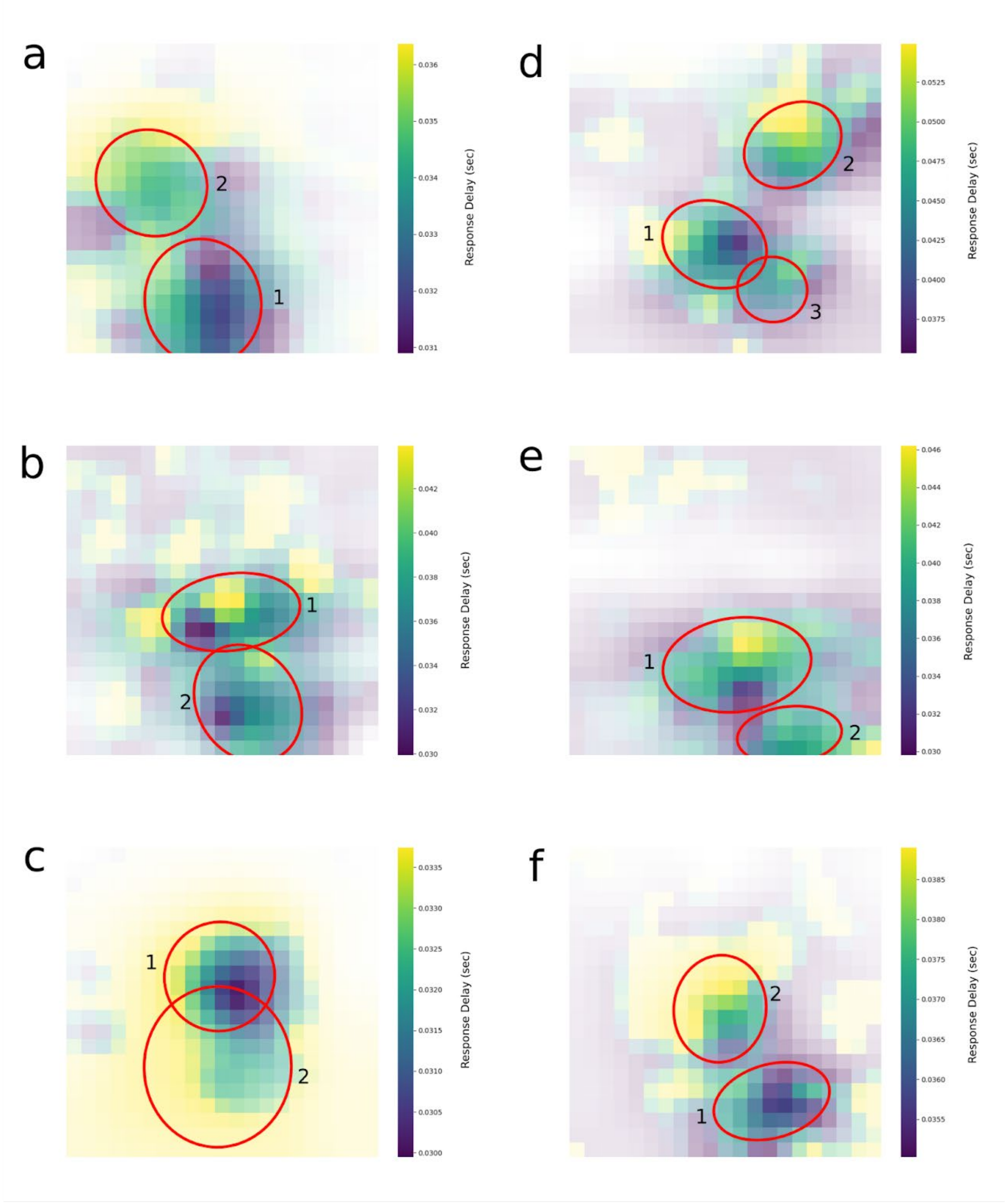

**Supplementary Fig. 2. Heatmaps of coupled photoreceptors' response delays.** (a-c) Apis coupled photoreceptors' response delays. (d-f) Bombus coupled photoreceptors' response delays. Primary RFs (larger weighting) exhibit, in general, shorter response delays than secondary ones. The only exception appears to be (c), for which the weightings of the two 2D Gaussian RFs were 0.51 (main RF) and 0.49 (secondary RF). For each heatmap 'cell', the delay to peak fitted response is displayed (as shown in Supplementary Fig. 1). Opacity of the heatmaps is scaled by relative sensitivity. Red circles correspond to the FWHM of the fitted 2D-Gaussian RFs (as in Fig. 1c,e).

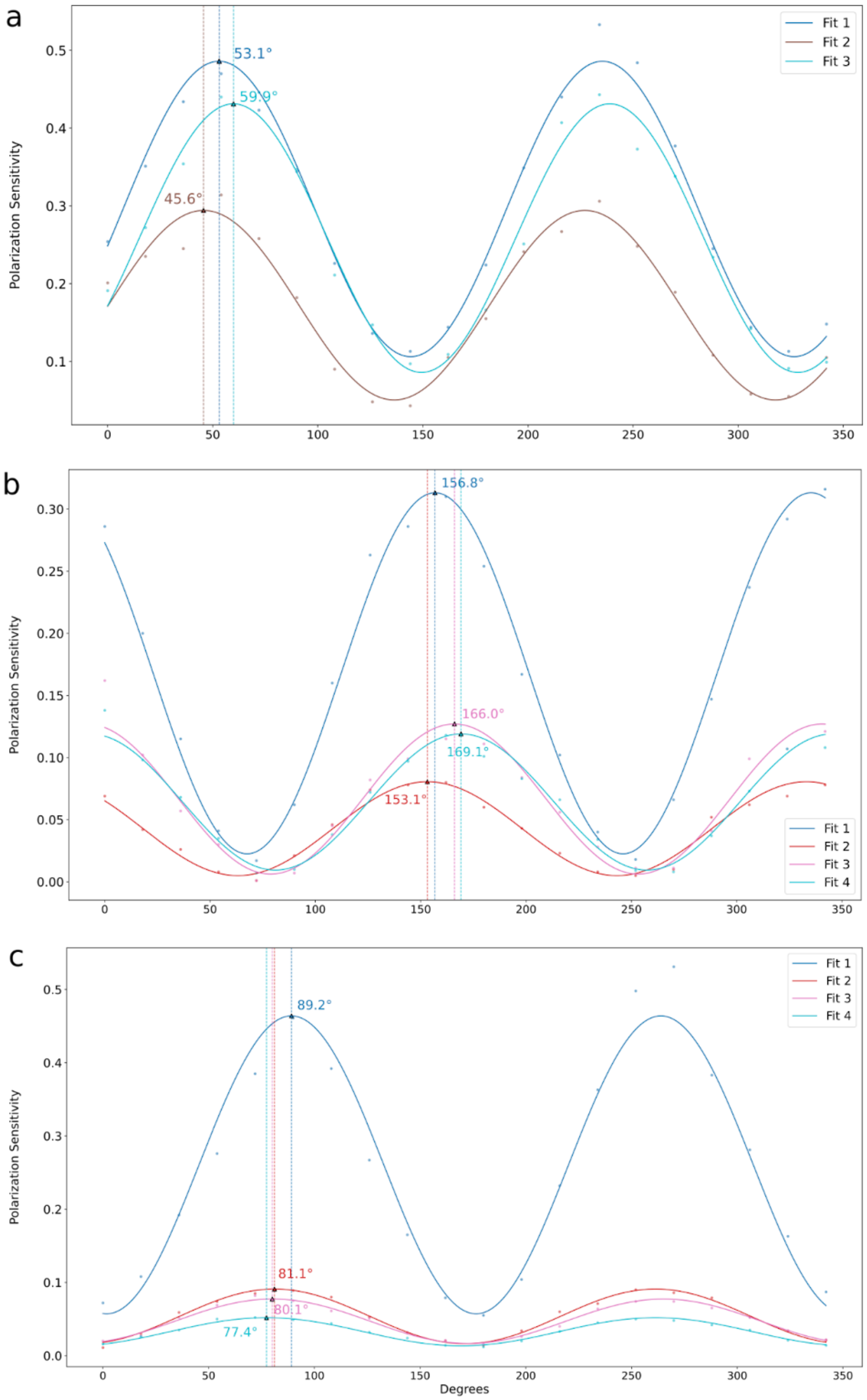
a
b
c
Polarization Sensitivity
Degrees
Fit 1
Fit 2
Fit 3
Fit 4
53.1°
59.9°
45.6°
156.8°
166.0°
169.1°
153.1°
89.2°
81.1°
80.1°
77.4°

**Supplementary Fig. 3. (a-c) $\cos^2$ fits of polarization sensitivity data and $\varphi_{max}$ shifts of coupled DRA cells in three different coupled cells.** In each panel, each curve corresponds to the polarization sensitivity of a coupled cell. The $\varphi_{max}$ values are annotated at the maxima of the curves. Shifts in $\varphi_{max}$ angles provide evidence that these DRA cells are indeed coupled with neighbouring cells with slightly different $\varphi_{max}$ angles.

**Supplementary Table 1. Model selection for receptive field size (Full Width at Half Maximum; FWHM) ~ polarization sensitivity (PS).** (a) Models with reduced interaction terms were compared against the full model and the best model was chosen in terms of the Akaike Information Criterion (AIC; in red). (b) Nested models were compared against the best model and the best fit was chosen in terms of the Likelihood-ratio test and the AIC (in red). (c) The model chosen after model selection. The increase of the FWHM is an additive function of polarization sensitivity, eye region and species.

| **Model (FWHM~)** | **AIC** | **ΔDeviance** | **Δd.f.** | **p-value** |
|---|---|---|---|---|
| *a. Compared against full model (PS * region * species)* | 142.96 | | | |
| PS:region:species | 139.29 | 0.3306 | 2 | 0.8476 |
| region:species | 136.64 | 1.6731 | 4 | 0.7956 |
| PS:species | 134.68 | 1.7153 | 5 | 0.887 |
| species | 139.33 | 8.3658 | 6 | 0.2125 |
| PS:region | 137.64 | 10.6783 | 8 | 0.2206 |
| *b. Compared against best model based on AIC (PS + region + species + PS:region)* | | | | |
| PS:species | 135.59 | 4.9149 | 2 | 0.0857 |
| species | 139.33 | 6.6505 | 1 | 0.0099 |
| region | 155.81 | 29.1301 | 4 | 0 |
| *c. Chosen model* | | | | |
| FWHM ~ PS + region + species | | | | |

#### References

1. J. Zhou, N. C. Benson, K. Kay, J. Winawer, Predicting neuronal dynamics with a delayed gain control model. *PLoS Comput. Biol.* **15**, e1007484 (2019).
